# Use of eVLP-based vaccine candidates to broaden immunity against SARS-CoV-2 variants

**DOI:** 10.1101/2021.09.28.462109

**Authors:** Jasminka Bozic, Tanvir Ahmed, Barthelemy Ontsouka, Anne-Catherine Fluckiger, Abebaw Diress, Tamara Berthoud, Xiaoyang Yuan, Lanjian Yang, Francisco Diaz-Mitoma, David E. Anderson, Catalina Soare

**Affiliations:** VBI Vaccines, 201-310 Hunt Club Road, Ottawa ON K1V 1C1, Canada; Bionaria, 11Bis Rue de la Garenne, 69290 Saint-Genis-les-Ollières, France; VBI Vaccines, Cambridge, 222 Third Street, Cambridge, 02142 Massachusetts, USA

**Keywords:** SARS-COV-2 variants, Vaccine, Virus-like-particles, Immunogenicity, cross-neutralizing antibodies

## Abstract

Rapid emergence of SARS-CoV-2 variants is a constant threat and a major hurdle to reach heard immunity. We produced VBI-2905a, an enveloped virus-like particle (eVLP)-based vaccine candidate expressing prefusion spike protein from the Beta variant that contains several escape mutations. VBI-2905a protected hamsters against infection with a Beta variant virus and induced high levels of neutralizing antibodies against Beta RBD. In a heterologous vaccination regimen, a single injection of VBI-2905a in animals previously immunized with VBI-2902, a vaccine candidate expressing S from ancestral SARS-CoV-2, hamsters were equally protected against Beta variant infection. As an alternate strategy to broaden immunity, we produced a trivalent vaccine expressing the prefusion spike protein from SARS-CoV-2 together with unmodifed S from SARS-CoV-1 and MERS-CoV. Relative to immunity induced against the ancestral strain, the trivalent vaccine VBI-2901a induced higher and more consistent antibody binding and neutralizing responses against a panel of variants including Beta, Delta, Kappa, and Lambda, with evidence for broadening of immunity rather than just boosting cross-reactive antibodies.

## Introduction

The outbreak of a severe respiratory disease in Wuhan, China in December 2019 led to the identification of a new betacoronavirus related to the severe acute respiratory syndrome (SARS) coronavirus that was named SARS-CoV-2 (Wu et al., 2020). SARS-CoV-2 rapidly spread worldwide in a global pandemic in 2019 (COVID-19) and was declared a public health emergency of international concern (World Health Organization, 2020). Unprecedented effort and innovation in vaccine development resulted in vaccines, deployed under emergency use authorization, against SARS-CoV-2 in less than a year (FDA, 2020).

Coronaviruses are large single-strand RNA viruses with replication that is error-prone despite some proofreading mechanisms (Smith and Denison, 2012). Resulting mutational changes can either be detrimental and lead to viral extinction or confer advantage to the virus and result in better adaptation to the host. Massive replication of SARS-CoV-2 on a global scale contributes to increasing numbers of mutations and emergence of variants. Variants of concern (VOC) are defined by clear evidence indicating a significant impact on transmissibility, severity, and/or immunity that is likely to have an impact on the disease epidemiology (EDCC, 2021). In July 2021, four VOC that first emerged as locally dominant variants before spreading globally were discovered in the U.K. (B.1.1.7 – “Alpha”), South Africa (B.1.351 – “Beta”), Brazil (P1 – “Gamma”), and more recently in India (B.1.617 – “Delta”). In late August the Mu variant first emerged in Columbia, joining this list of Variants of Interest (VOI) together with Lambda while the incidence of Alpha was decreasing.

The CoV spike (S) protein contains a receptor binding domain (RBD) critical for binding to and infection of host cells, and is a major target for mutational changes which enhance adaptation to the host (Berrio et al., 2020; Boni et al., 2020). Each of the VOC are characterized by a number of shared mutations expressed on S, primarily located in the RBD and N-terminal domain (NTD), that serve to increase inter-individual transmission, escape neutralizing antibodies acquired by vaccination or prior natural SARS-CoV-2 infection, or both (Harvey et al., 2021; Zhang et al., 2020). For instance, the first identified VOC, Alpha, is characterized by a D614G mutation, among other mutations, that is now fixed in all globally circulating variants of the virus. D614G is associated with increased transmissibility (Plante et al., 2021; Zhang et al., 2020) but does not have a major impact on neutralization by serum from either vaccinated or COVID-19 convalescent individuals. The following VOC that emerged in South Africa was rapidly identified as a vaccine-escape mutant. This Beta variant bears several mutations in its RBD, including E484K and N501Y, which significantly inhibit neutralizing activity elicited against the Ancestral Wu-1 strain of virus whether acquired by vaccination or infection (Tegally et al., 2020; Cele et al., 2021). Emergence of escape mutants is a major concern because most of the licensed vaccines are based on expression of various forms of S using the Ancestral sequence of the S protein (Hoffmann et al., 2021 ; Kyriakidis et al., 2021; Lamb, 2021). More recently, the Delta variant spread from India to many countries with great speed in spite of significant proportions of fully vaccinated invididuals in many countries. Delta shows the RBD mutation L452R which appeared independently in several areas of the globe, including in variants Lambda, Kappa, Epsilon, Iota, and contributes to escape neutralization from Abs induced by previously acquired immunity (Deng et al.2021). Additionally, mutation P681R in the furin cleavage site of Delta could increase the rate of S1-S2 cleavage, resulting in better transmissibility (Cherian et al., 2021).

Recently, we developed a SARS-CoV-2 candidate vaccine, VBI-2902a, comprised of enveloped virus-like particles (eVLPs) expressing a modified prefusion form of the ancestral S sequence, adjuvanted with aluminum phosphate (Alum). We recently demonstrated that VBI-2902a induced strong neutralizing activity in mouse immunogenicity studies, and protected hamsters from SARS-CoV-2 challenge using a virus related to the ancestral isolate (Fluckiger et al. 2021). Interim results from a Phase I clinical study in healthy, seronegative individuals (ClinicalTrials.gov Identifier: NCT04773665) demonstrated robust (4.3-fold greater) neutralizing activity 28 days after a second, 5µg dose of VBI-2902a, relative to a panel of COVID-19 convalescent sera.

Employing the same strategy, we produced a new vaccine candidate, VBI-2905a, that expresses a modified prefusion S based on the Beta variant sequence. Consistent with previous studies, VBI-2905a elicited neutralizing antibody responses against the Beta variant which were significantly greater than those induced by VBI-2902a, and responses against the ancestral strain which were comparable to VBI-2902a. Consistent with the role of neutralizing antibody responses as a presumed correlate of protection, greater efficacy was observed in hamsters vaccinated with VBI-2905a relative to VBI-2902a when challenged with the Beta variant. Noteworthy was the observation in an alternative vaccination regimen that priming with VBI-2902a followed by a single booster dose of VBI-2905a induced strong neutralizing antibody responses against the ancestral strain as well as both Beta and Delta VOC.

We also evaluated immunity elicited with a distinct eVLP-based candidate, VBI-2901a, which expresses a modified prefusion S based on the ancestral sequence in addition to the related S proteins from SARS CoV-1 and MERS. Immunization with VBI-2901a induced neutralizing antibody titers against the Beta variant significantly greater than VBI-2902a and comparable to those induced with VBI-2905a, effectively broadening immunity to VOC not contained within the vaccine. Antibody binding and neutralizing titers against an extended panel of variants demonstrated responses typically 3-fold greater than that observed with VBI-2902a. Collectively, these results demonstrate multiple ways to broaden immunity to SARS-CoV-2 VOC.

## Material and Methods

### Plasmids, eVLP production, and adjuvant formulation

Expression plasmid for the production of eVLPs expressing SARS-CoV-2 S proteins have been described previously (Fluckiger et al. 2021). Briefly, the prefusion modified form of S was obtained by introducing a mutation at the furin cleavage site (RRAR → GSAS) and two Proline at position K986-V987 of the Wuhan reference and swapping the transmembrane cytoplasmic domain with that of the VSV-G protein. VBI-2902a was produced using the Wuhan-Hu-1 spike sequence (Genbank accession number MN908947), and VBI-2905a was produced using the same strategy with S sequence from Beta variant B.1.351 isolate EPI_ISL_911433 (GISAID). Production and purification of eVLPs were conducted as described elsewhere (Fluckiger et al. 2021). The preparation of eVLPs expressing either Wuhan reference Spike or Beta variant Spike were formulated in Aluminum phosphate (Alum, Adjuphos^®^, Invitrogen) to obtain vaccine candidate VBI-2902a and VBI-2905a, respectively. To produce VBI-2901, Two additionnal plasmids were produced that expressed the optimized sequences for full-lenght unmodified S protein from SARS-CoV-1 and MERS-CoV. To produce trivalent eVLPs, HEK-293SF-3F6 were cotransfected with these 2 plasmids together with the plasmid coding for prefusion ancestral SARS-CoV-2 S used for VBI-2902a production, and the MLVGAG plasmid as described. Expression of SARS-CoV-2 S, SARS-CoV-1 S, MERS-CoV S and GAG were determined by Western blot analysis (Supplementary material Fig.S1).

### Mouse immunization study

Six- to 8-week-old female C57BL/6 mice were purchased from Charles River (St Constant, Quebec Canada). The animals were acclimatized for a period of at least 7 days before any procedures were performed. The animal studies were conducted under ethics protocols approved by the NRC Animal Care Committee. Mice were maintained in a controlled environment in accordance with the “Guide for the Care and Use of Laboratory Animals” at the Animal Research facility of the NRC’s Human Health Therapeutics Research Centre (Montreal). Mice were randomly assigned to experimental groups of 10 to 15 mice and received intraperitoneal (IP) injections with 0.5 mL of adjuvanted SARS-CoV-2 eVLPs as described elsewhere (Fluckiger, 2021). Blood was collected on day -1 before injection and day 14 after each injection for humoral immunity assessment at time of euthanasia.

### Hamster challenge study

Syrian golden hamsters (males, 5-6 weeks old) were purchased from Charles River Laboratories (Saint-Constant, Quebec, Canada). The study was conducted under approval of the CCAC committee at the Vaccine and Infectious Disease Organization (VIDO) International Vaccine Centre (Saskatchewan, Canada). Animals were randomly assigned to each experimental group (A, B) (n=10/group). Animals received 2 intramuscular (IM) injection of either 0.9%-saline buffer (saline control group) or VBI-2902a (VBI-2902a group), or VBI-2905a (VBI-2905a group), or a first dose of VBI-2902a followed by a second injection of VBI-2905a (Heterologous boost group). Each dose of eVLP-based vaccine contained 1µg of Spike protein formulated with 125 µg of Alum. Injection was performed by intramuscular (IM) route at one side of the thighs in a 100 µL volume. The schedule for immunization, challenge and sample collection is depicted on Fig. 2a. All animals were challenged intranasally via both nares with 50 μL/nare containing 1×10^5^ TCID50 of hCoV-19/South Africa/KRISP-EC-K005321/2020 (Seq. available at GISAID: EPI_ISL_678470) strain per animal. Body weights and body temperature were measured at immunization for 3 days and daily from the challenge day. General health conditions were observed daily through the entire study period. Blood samples were collected as indicated on Fig. 2a.

### Antibody binding titers

Anti-SARS-CoV-2 specific IgG binding titers in sera were measured by standard ELISA procedure described elsewhere (Kirchmeier et al., 2014), using recombinant SARS-CoV-2 S RBD proteins (Sinobiological). For total IgG binding titers, detection was performed using a goat anti-mouse IgG-Fc HRP (Bethyl), or Goat anti-Hamster IgG HRP (ThermoFisher), or goat anti-human IgG heavy and light chain HRP-conjugated (Bethyl). HRP-conjugated Goat anti-mouse IgG1 and HRP-conjugated goat anti-mouse IgG2b HRP (Bethyl) were used for the detection of isotype subtype. Determination of Ab binding titers to Spike RBDs was performed using SARS-COV-2 RDB recombinant protein for the specificty of choice as described in Suppl. Table 1. The detection was completed by adding 3,3’,5,5’-tetramethylbenzidine (TMB) substrate solution, and the reaction stopped by adding liquid stop solution for TMB substrate. Absorbance was read at 450 nm in an ELISA microwell plate reader. Data fitting and analysis were performed with SoftMaxPro 5, using a four-parameter fitting algorithm.

### Virus neutralization assays

Neutralizing activity in mouse serum samples was measured by standard plaque reduction neutralization test (PRNT) on Vero cells at the NRC using 100 PFU of SARS-CoV-2/Canada/ON/VIDO-01/2020 (Wu-1 virus) or hCoV-19/South Africa/KRISP-EC-K005321/2020 (Beta virus). Results were represented as PRNT90 end point titer (EPT), corresponding to the lowest dilution inhibiting respectively 90% of plaque formation in Vero cell culture.

### Neutralization assay with pseudoparticles

Production of pseudoparticles (pp) pseudotyped with various spike proteins and neutralization assay was adapted from Dreux et al, 2009. Expression plasmid were designed using full length S protein sequences as described previsouly. Accession number and mutations are listed in Suppl. Material Table 1. We produced infectious SARS-CoV-2pp carrying a GFP-firefly luciferase double reporter gene (plasmid pjm155, Garrone et al., 2011) instead of green fluorescent protein (GFP). Luciferase activity in infected hACE2-HEK293 cells was measured with a Bright-Glo Luciferase assay system (Promega) and a Beckman Coulter DTX880 plate reader. Data were expressed in relative luminescence units (RLUs). The percentage of neutralization was calculated by comparing the luciferase activity in cells infected with SARS-CoV-2pp in the presence of serum from immunized animals with luciferase activity in cells infected with SARS-CoV-2pp in the absence of serum.

### Statistics

All statistical analyses were performed using GraphPad Prism 9 software (La Jolla, CA). Unless indicated, multiple comparison was done with Dunn’s corrected Kruskall-Wallis test on unpaired samples and Friedman test on paired samples. The data were considered significant if p < 0.05. Geometric mean titers (GMT) with standard deviation are represented on graphs. No samples or animals were excluded from the analysis. Randomization was performed for the animal studies.

## Results

### Heterologous boosting with eVLPs bearing S protein from the Beta variant broadens immunity

VBI-2902a is an eVLP-based vaccine candidate that expresses a modified prefusion SARS-CoV-2 S protein from the ancestral Wu-1 strain, adjuvanted with Alum (Fluckiger et al., 2021). VBI-2905a expresses a modified prefusion SARS-CoV-2 S protein from the Beta variant and is also adjuvanted with Alum. We immunized mice with 2 injections of VBI-2902a, 2 injections of VBI-2905a, or a first injection of VBI-2902a followed by a second injection of VBI-2905a (heterologous boost). As previously described (Fluckiger et al., 2021), 2 doses of VBI-2902a induced high levels of neutralizing Ab response against the ancestral Wu-1 strain (GMT = 2,458) which were significantly reduced against the Beta variant (GMT = 94) (Fig.1a-b). By contrast, VBI-2905a induced Abs that neutralized Beta and ancestral viruses at similar levels in mice, yielding only a 2.2-fold difference with non significant *p* = 0.1484 (Fig. 1a-b). Sera from mice in the heterologous boost group cross-neutralized both the Beta variant and the ancestral strain with similar potencies (1,4 fold difference with *p =* 0.3828). Heterologous boosting with VBI-2905a significantly increased the PRNT90 against the ancestral strain compared to 2 doses of VBI-2905a alone (from GMT of 371 to 820, *p* = 0.0267) to levels that were closer to those reached after two doses of VBI-2902a (*p* = 0.0131), while PRNT90 GMTs against the Beta variant were comparable to 2 doses of VBI-2905a (respectively GMT = 564 vs GMT = 619, *p* = 0.8785).

**Figure 1:**
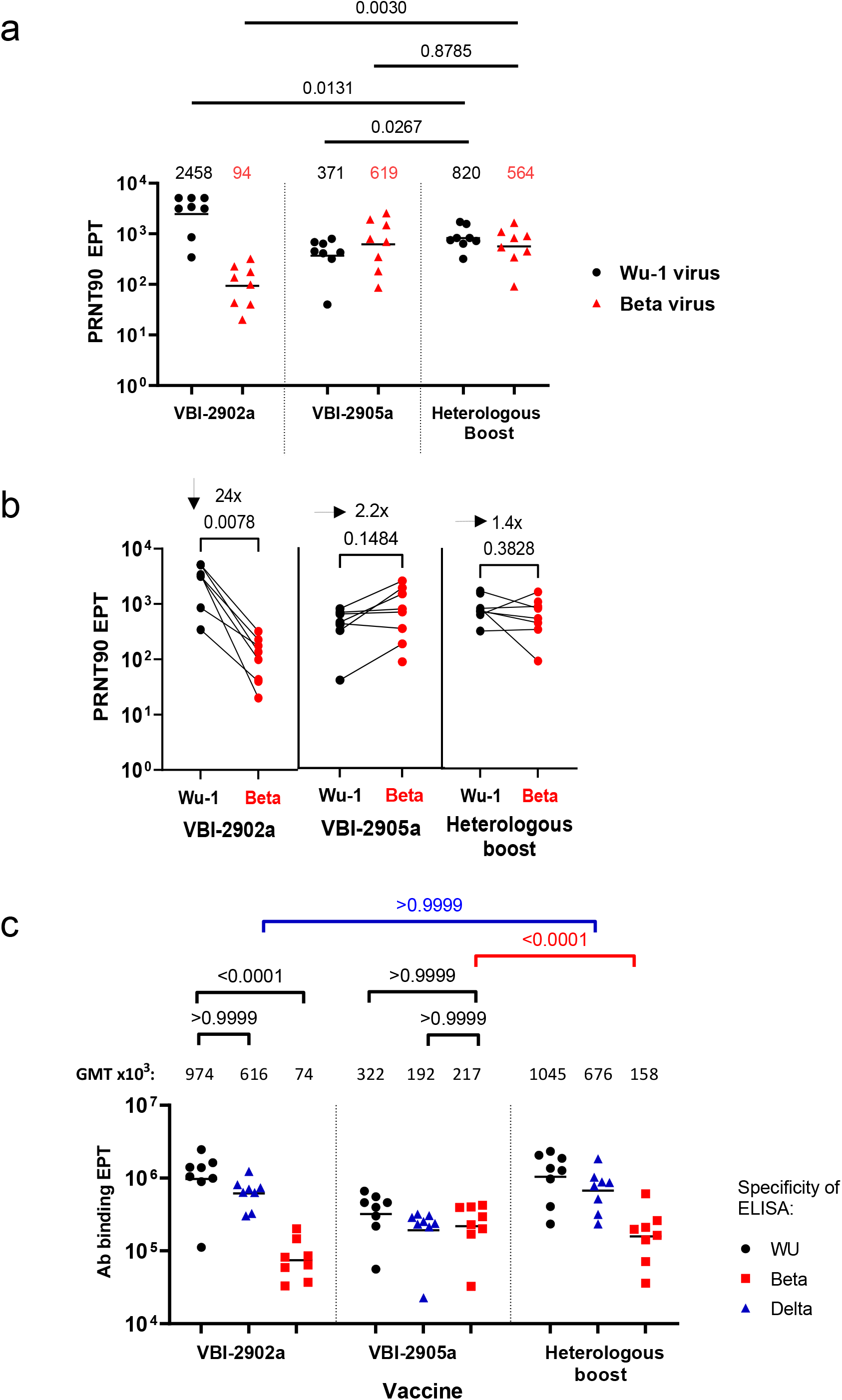
Immunogenicity of VBI-2902a and VBI-2905a in mice. C57BL/6 mice, 8 per group, received 2 IP injections 3 weeks apart, of VBI-2902a or VBI-2905a or a first injection of VBI-2902a followed by a second injection of VBI-2905a (Heterologous boost), each containing 0.1 µg of S. Blood was collected at day 14 after the second injection for monitoring of the humoral response. **(a)** Sera from each group were analyzed in PRNT assay with a 90% threshold (PRNT90) using Wu-1 virus and Beta virus as described in Material and Methods. GMT and results from two-tailed Mann–Whitney U-test are indicated. **(b)** Change in neutralization between Wu-1 and Beta viruses. Fold change was calculated for each serum as the ratio between reactivity to Ancestral and Beta RBD, Fold change in each group is indicated as the geometric mean preceeded by an arrow. Statistical analysis was determined using two tailed Wilcoxon test. **(c)** Ab binding titers were evaluated by ELISA using recombinant Delta RBD as described in Material and Methods. Statistical significance was determined by Kruskall-Wallis test.

Analysis of Ab binding titers to S protein RBDs was consistent with the neutralization data (Fig.1c). VBI-2902a induced high levels (most of the sera >10^6^ EPT with GMT 974×10^3^) of Ab binding titers against the ancestral S RBD with significantly reduced cross-reactivity against the Beta variant RBD (GMT 74×10^3^), though there was good cross-reactivity against the Delta variant RBD (GMT 616×10^3^). Antisera from immunization with VBI-2905a showed similar crossreactivty against Ancestral, Delta and Beta RBD (respectively GMT 322×10^3^, 192×10^3^ and 217×10^3^). Animals receiving the heterologous prime boost regimen had similar reactivity to ancestral and Delta RBD as compared with the VBI-2902a group, and similar reactivity to Beta RBD compared to the VBI-2905a group.

### Heterologous boosting with VBI-2905a protects hamsters against SARS-COV-2 Beta variant

Golden Syrian hamsters were intramuscularly vaccinated 3 weeks apart with two doses of eVLP vaccine candidates, comprised of: two doses of VBI-2902a (group VBI-2902a), two doses of VBI-2905a (group VBI-2905a), or a priming dose of VBI-2902a followed by a second, booster dose of VBI-2905a (group heterologous boost) (Fig. 2a).

**Figure 2:**
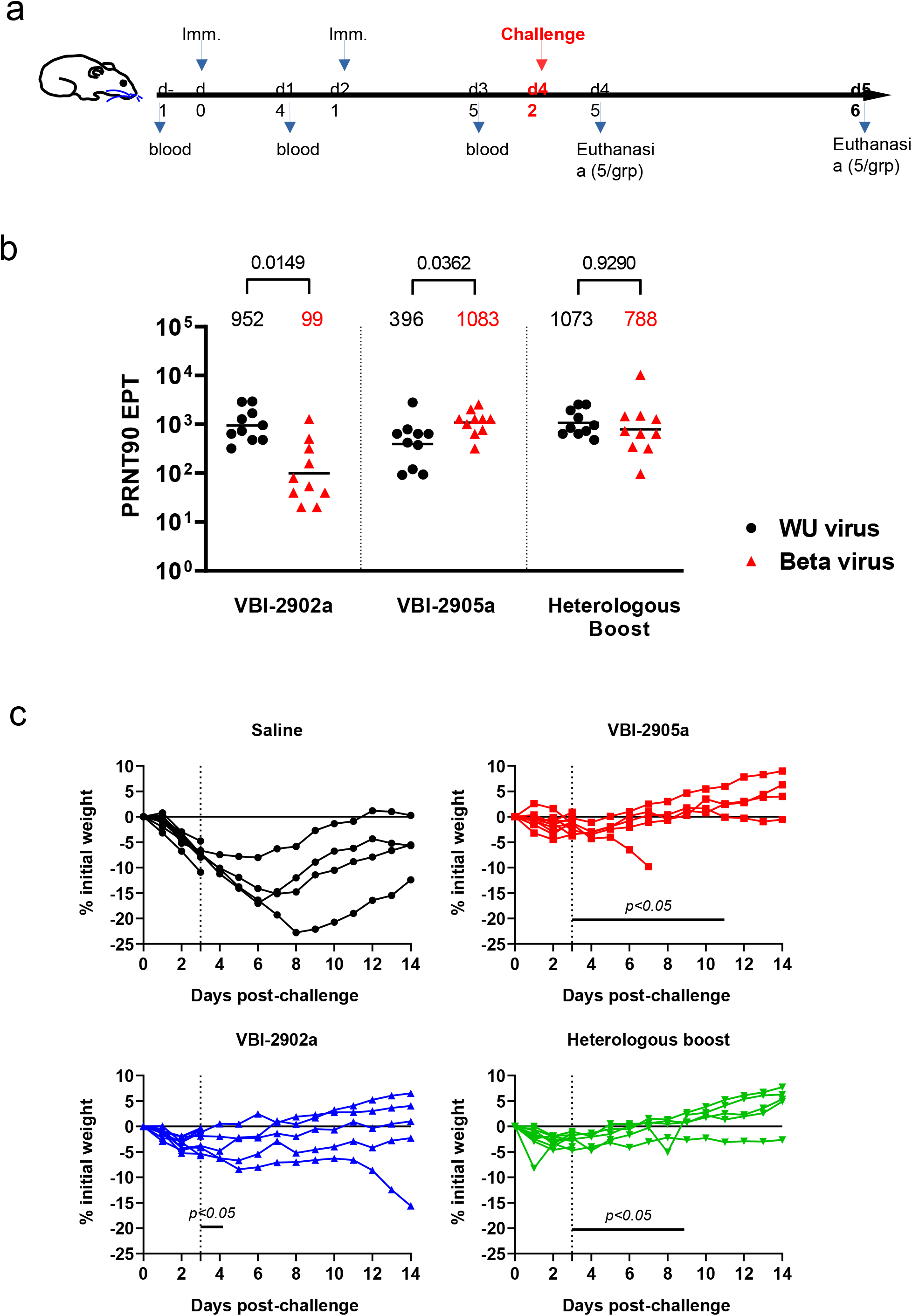
Beta variant challenge in Syrian golden hamsters after immunization with VBI-2902a and VBI-2905a. **(a)** Schematic representation of the challenge experiments. Four groups of 10 Syrian gold hamsters received 2 IM injections 3 weeks apart, of placebo saline buffer or VBI-2902a or VBI-2905a or a first injection of VBI-2902a followed by a second injection of VBI-2905a, with 1µg of S per dose. Animals in Placebo groups received Saline buffer. Blood was collected 2 weeks after each injection. Three weeks after the last injection (day 42) hamsters were exposed to SARS-CoV-2 Beta virus at 1×10^5^ TCID50 per animal via both nares. At 3 days post infection (dpi), 5 animals per groups were sacrificed for viral load analysis. The remaining animals were clinically evaluated daily until end of study at 14dpi. **(b)** Neutralization activity was measured by PRNT90 in immunized groups; results are represented as PRNT90 EPT. GMT and statistical significance from two tailed Friedman test are indicated **(c)** Hamsters were monitored daily for weight change. Results are represented for each animal in each groups as kinetic of weight change from day 0 to day 14 after infection. One animal from VBI-2905a group was sacrificed at day 7 because of worsening of clinical presentation after a fight in the cage. Significant days of weight loss relative to Saline group (p<0.005) are indicated. Statistical analysis was performed with unpaired non parametric multiple t test using Holm-Šidák method.

Neutralizing activities titers against the ancestral virus were comparable across all groups, including hamsters immunized with 2 doses of the Beta S candidate (VBI-2905a) (Fig.2b). Neutralization of the Beta variant was lower after immunization with VBI-2902a, with a significant 9.6-fold decrease of Beta nAb compared to homotypic immunization with VBI-2905a (GMT 99 in VBI-2902a and 1083 in VBI-2905a, p = 0,0033). In contrast, nAb titers against Beta RBD were similar in groups that received either two doses of VBI-2905a or heterologous boosting.

Three weeks after the second immunization, hamsters were exposed to 1×10^5^ TCID50 of the Beta variant virus in each nare. In the placebo group, hamsters began losing weight the day after infection which continued until day 6-8. Vaccination with 2 doses of VBI-2902a based on the ancestral S protein induced limited protection against challenge with moderate weight loss recorded until day 4, and only a fraction (3/5) of the animals fully regained their initial body weight after day 7. By contrast, hamsters vaccinated with 2 doses of VBI-2905a exhibited transient weight loss up to day 2-3 and then rapidly regained weight. A similar pattern was observed in hamsters that received VBI-2905a as a boost. As we have observed in previous hamster challenge studies of VBI-2902a, there was a correlation between neutralizing antibody titers against the Beta variant and protection from disease (weight loss) after challenge (data not shown).

### Immunization with a pan-coronavirus candidate may protect against VOC not contained within the vaccine

We hypothesized that exposing the immune system to multiple spike proteins at the same time might help broaden humoral immunity that could recognize emerging variants or new coronaviruses more phylogenetically distant to the vaccine candidate. To test this hypothesis we produced VBI-2901a, a trivalent eVLP vaccine formulated with Alum, that expresses a prefusion form of the ancestral SARS-CoV-2 S (identical to VBI-2902) with unmodified full length S from SARS-CoV-1 and MERS-CoV (Suppl. Fig. S1). Mice that received 2 doses of trivalent VBI-2901a had increased nAb titers (GMT 2915) against the ancestral virus relative to mice that received 2 doses of monovalent VBI-2902a (GMT 831) or VBI-2905a (GMT 448) (Fig. 3a). Moreover, trivalent VBI-2901a induced neutralization activity against the Beta virus that was equivalent to what was observed in response to homotypic VBI-2905a, and significantly higher than that observed after VBI-2902a vaccination (Fig. 3a). Neutralization of both Delta and Kappa variant pseudotyped particles confirmed broadened neutralizing immunity elicited by VBI-2901a, with titers approximately 3-fold greater than those induced by VBI-2902a (Fig. 3b). Consistent with the neutralization activity, VBI-2901a induced higher and/or more consistent levels of Ab binding to the RBD among all variants evaluated, including Beta, Delta, and Lambda (Fig. 3c).

**Figure 3:**
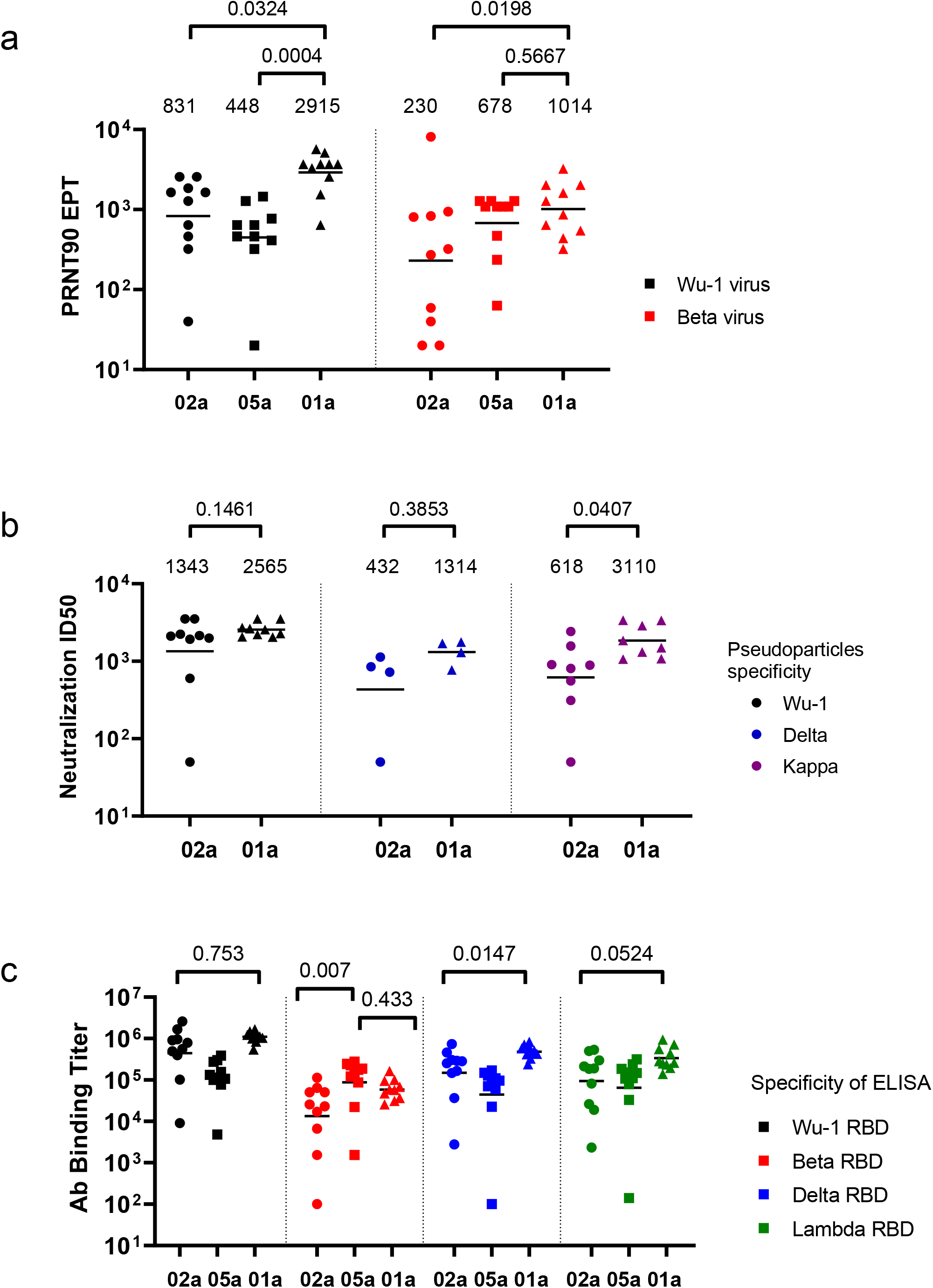
Immunogenicity of trivalent VBI-2901a. Three groups of 10 mice were immunized with 2 doses of VBI-2901a (01a) or VBI-2902a (02a) or VBI-2905a (05a) 3 weeks apart. Blood was collected at day 14 after the last injection for monitoring of the humoral response. **(a)** Neutralization EPT measured by PRNT90 against Wu-1 virus and Beta variant. GMT are indicated above each group. **(b)** Neutralization of pseudoparticles expressing S from Wu-1 virus, or Delta or Kappa variants are represented as half-maximum inhibitory dilutions (Neutralization ID50). Geometric means are indicated above each panel. Due to technical limitations, only 8 sera per groups were tested against Wu-1 and Kappa pseudoparticles and 4 sera against Delta pseudoparticles. Sera were randomly picked. **(c)** Ab binding titers measured in ELISA against recombinant RBD from Wu-1 ancestral virus, or Beta, Delta, and Kappa variants.

## Discussion

Less than a year after identification of the new SARS-CoV-2 virus, variants emerged with impact on transmissibility, severity and immunity (EDCC, 2021) that challenge the development and durability of vaccine strategies designed to reach herd immunity. Indeed, all approved vaccines have been designed against the ancestral SARS-COV-2 virus that is no longer circulating but has been replaced by variants containing mutations which are enabling escape from nAbs induced against the ancestral strain (Berio et al. 2020; Boni et al, 2020). In the present study, we compared several strategies to broaden antibody-based immunity which is presumed to be a correlate of protection against SARS-CoV-2.

In addition to our eVLP vaccine expressing the prefusion S from the ancestral SARS-CoV-2 virus, we produced an eVLP-based vaccine expressing the prefusion S from Beta variant. Beta was chosen for its deleterious mutations E484K and K417N, which enable escape neutralization from ancestral virus mAbs (Hoffman et al, 2021). We have previously demonstrated that 2 doses of VBI-2902a protected hamsters against infection by the ancestral Wuhan SARS-CoV-2 virus and we confirmed here that VBI-2905a also protected hamsters from infection with the SARS-CoV-2 Beta variant. We have also demonstrated that a heterologous boost with Beta variant vaccine VBI-2905a given to animals that had received a single priming dose of ancestral strain vaccine VBI-2902a protected against the new Beta variant while also maintaining cross-reactivity against the ancestral strain Moreover, heterologous eVLP boosting with VBI-2905a also induced high levels of antibody reactivity against the globally dominant Delta VOC. Additional challenge studies are in progress to evaluate if a heterologous boosting strategy can confer protection in Syrian golden hamsters against infection with the Delta variant.

Building upon the flexibility of the eVLP vaccine technology, we produced VBI-2901a, a multivalent coronavirus candidate containing S proteins from SARS-CoV-2, SARS-CoV, and MERS-CoV with the intent to broaden immunity to emerging VOC as well as novel, related betacoronaviruses that may infect humans in the future. Vaccines currently in use or in clinical evaluation that are based on the ancestral strain induce neutralizing antibody responses that are less reactive against the Beta VOC, with titers typically 5-10 lower than against the ancestral strain (Wibmer et al., 2021; Wang et al., 2021). In marked contrast, VBI-2901a elicited robust nAb responses not only against the ancestral SARS-CoV-2 strain, but also against the Beta variant, providing evidence of the vaccine candidate’s ability to broaden immunity and “anticipate” an emerging variant not contained within the vaccine. High levels of cross-neutralizing activity elicited by VBI-2901a were also observed against the Delta and Kappa variants. Other studies have shown that plasma from individuals previously infected with SARS-CoV-1 who received the BNT162b2 mRNA vaccine, which is based on the ancestral SARS-CoV-2 virus, contained a broad spectrum of neutralizing antibodies against 10 sarbecoviruses tested, including SARS-COV-2 variants, several strains of SARS-CoV-1, and Bat and Pangolin CoV (Tan et al., 2021). Further studies are underway to better understand how VBI-2901a, which similarly exposes the B cell repertoire to spike proteins from both SARS-CoV-1 and SARS-CoV-2, broadens neutralizing activity against SARS-CoV-2 variants as well as to assess neutralizing responses to phylogenetically more distant coronaviruses.

Broadening of the neutralizing antibody response has also been shown using nanoparticles of mosaïc RBD from various betacoronavirus species (Cohen et al., 2021; Walls et al., 2021). However, the N terminal domain of the S protein is another important target for neutralizing antibodies and the site of many mutations that could potentially contribute to antibody neutralization escape (Andreoni et al. 2021). Given that VBI-2901a expresses the full-length ectodomain of the Coronaviruses spike, it will be critical to determine the respective roles and importance of the RBD, NTD, and the highly conserved S2 domains in broadening immunity.

Whereas vaccines based on the ancestral strain of SARS-CoV-2 protect against severe disease caused by variants of concern, variants such as Beta are less sensitive to vaccine-induced immunity and efficacy rates are accordingly lower. This is likely to become more apparent as vaccine-induced immunity wanes and as variants continue to emerge with even greater numbers of mutations. One strategy to address these concerns is to administer booster doses to increase neutralizing antibody titers against the ancestral strain, a subset of which may cross-neutralize variants of concern. We have described three alternate strategies that have the potential to broaden immunity to a greater extent. An eVLP-based candidate based on the Beta variant S protein, VBI-2905a, induces potent immunity against not just the Beta virus, but also against the ancestral strain, though it is less potent against the Delta variant. However, building upon immunity induced against the ancestral strain with a priming dose of VBI-2902a, a single booster dose of VBI-2905a resulted in potent and more balanced neutralizing antibody responses against the ancestral virus, and Beta and Delta variants. Finally, we have described a novel trivalent eVLP candidate, VBI-2901a, which elicited potent and broad immunity against all variants tested, including Beta, Delta, Lambda, and Kappa, with the testing for the potential to neutralize more distantly related viruses currently underway.

## Abbreviations

eVLP: enveloped virus-like particules
CoV: coronavirus
VOC: Variant of concern
VOI: varaint of interest
RBD: receptor binding domain
NTD: N-terminal domain
Ab: antibody
nAb: neutralizing antibody
MLV: murine leukemia virus
ELISA: enzyme-linked-immuno-sorbent-assay
PRNT: plaque reduction neutralization test
EPT: end-point titer
Alum: aluminum phosphate
IP: IntraPeritoneal
IM: IntraMuscular
NRC: National Research Council Canada
VIDO: Vaccine and Infectious Disease Organization

## Acknowledgment

The authors want to thank Adam Asselin, Matthew Yorke, Teresa Daoud, Rebecca Wang, Gillian Lampkin (VBI vaccines) for outstanding technical support; NRC Animal Resources Group and the VIDO Saskatchewan team for remarkable care with animal experiments.

