## Supplementary Table S1 for "Use of eVLP-based vaccine candidates to broaden immunity against SARS-CoV-2 variants"

**Supplementary Table S1: Source of recombinant protein and S protein sequences used in the production of eVLPs**

| Recombinant protein | Manufacturer | Cat# | Description |
|---|---|---|---|
| SARS-CoV-2 Wu-1 ancestral | Sinobiological | 40589-V08B1 | SARS-CoV-2 (2019-nCoV) Spike Protein (S1+S2 ECD, His tag) |
| SARS-CoV-2 Wu-1 ancestral, RBD | Sinobiological | 40592-VNAH | SARS-CoV-2 (2019-nCoV) Spike Protein (RBD) |
| SARS-CoV-2 Beta variant RBD | Sinobiological | 40592-V08H85 | SARS-CoV-2 (2019-nCoV) Spike RBD (K417N, E484K, N501Y)-His |
| SARS-CoV-2 Delta variant RBD | Sinobiological | 40592-V02H3 | SARS-CoV-2 Spike RBD (L452R, T478K) Protein (Fc Tag) |
| SARS-CoV-2 Lambda variant RBD | Sinobiological | 40589-V08B23 | SARS-CoV-2 Spike S1+S2 (G75V, T76I, R246N, (247, 253) deletion, L452Q, F490S, D614G, T859N) Protein (ECD, His Tag) |
| SARS-CoV-2 Wu-1 S2 Subunit | Ray Biotech | 230-01103-100 | SARS-CoV-2 Spike Protein, S2 |
| SARS-CoV-1 S1+S2 | Sinobiological | 40634-V08B | SARS-CoV Spike S1+S2 ECD-His (S577A, Isolate Tor2) |
| MERS-CoV S1+S2 | Sinobiological | 40069-V08B | MERS-CoV Spike Protein (S1+S2 ECD, aa 1-1297, His Tag) |

| Genotype of S protein sequence | Source | Accession number |
|---|---|---|
| Wu-1 ancestral | Genbank | MN908947 |
| Beta variant isolate | GISEAD | EPI_ISL_3911433 |
| Delta variant isolate | GISEAD | EPI_ISL_2356230 |
| Lambda variant isolate | GISEAD | EPI_ISL_3368406 |
| Kappa variant isolate | GISEAD | EPI_ISL_1704611 |
| SARS-CoV-1 HKU-39849 | Genbank | JN854286.1 |
| MERS-CoV EMC/2012 | Genbank | JX869059.2 |
