## Supplementary Figure S1 for "Use of eVLP-based vaccine candidates to broaden immunity against SARS-CoV-2 variants"

### Supplementary Figure 1 : description of Trivalent VBI-2901

a

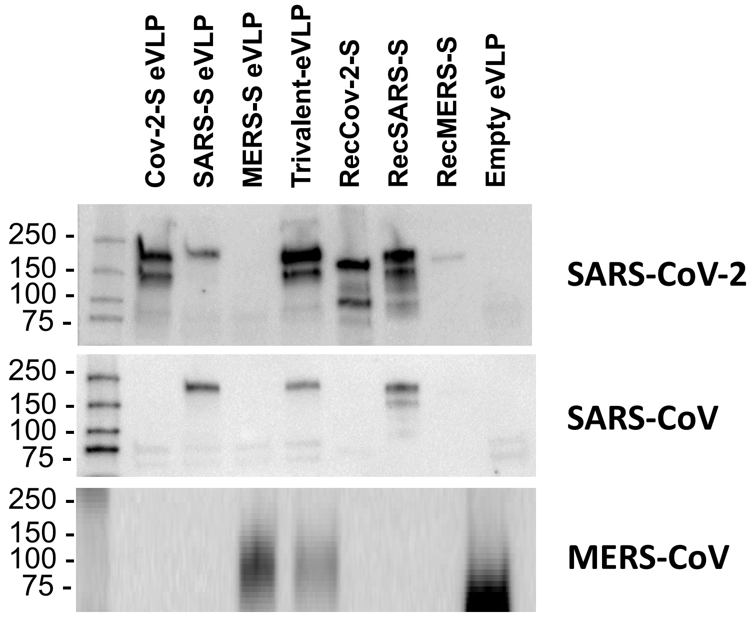


b

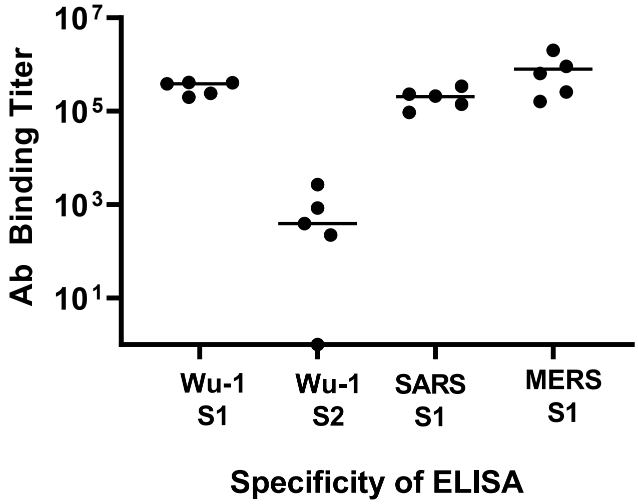


**Supplementary Figure S1: Production of trivalent eVLPs expressing S protein from 3 betacoronavirus species. (a)** Expression of S proteins analyzed by Western-blot in trivalent VBI-2901 recombinant proteins using Rabbit SARS-CoV-2 Spike RBD polyclonal Ab (Sinobiological) crossreacting with SARS-CoV-2 and SARS-CoV-1 S protein, or Rabbit MERS Coronavirus Spike protein S1 polyclonal Ab (Invitrogen). eVLPs produced with Gag plasmid only (empty eVLPs) and recombinant S proteins were used as negative and positive controls respectively. Each lane was loaded with purified eVLPs containing equivalent amount of 5µg of total protein evaluated by standard BCA assay **(b)** To test immunogenicity of trivalents eVLPs, 5 C57BL/6 mice received 2 doses of trivalent eVLPs formulated with Alum (VBI-2901a). Each dose contained 0.2 µg of S protein based on quantification by ELISA. Ab binding titers against Ancestral Wu-1 S1, Ancestral Wu-1 S2, SARS-CoV-1 S1 and MERS-CoV-1 S1 were measured by ELISA in sera collected 2 weeks after the last immunization using recombinant S protein as described in Suppl. table S1.
